# Concurrent evolution of anti-aging gene duplications and cellular phenotypes in long-lived turtles

**DOI:** 10.1101/2021.07.07.451454

**Authors:** Scott Glaberman, Stephanie E. Bulls, Juan Manuel Vazquez, Ylenia Chiari, Vincent J. Lynch

**Affiliations:** Department of Environmental Science and Policy, George Mason University, Fairfax, VA, USA; Department of Biology, University of South Alabama, Mobile, AL, USA; Department of Integrative Biology, University of California - Berkeley, Berkeley, CA, USA; Department of Biology, George Mason University, Fairfax, VA, USA; Department of Biological Sciences, University at Buffalo, SUNY, Buffalo, NY, USA

**Keywords:** turtles, aging, longevity, body size, cancer, ER stress, apoptosis

## Abstract

There are many costs associated with increased body size and longevity in animals, including the accumulation of genotoxic and cytotoxic damage that comes with having more cells and living longer. Yet, some species have overcome these barriers and have evolved remarkably large body sizes and long lifespans, sometimes within a narrow window of evolutionary time. Here, we demonstrate through phylogenetic comparative analysis that multiple turtle lineages, including Galapagos giant tortoises, concurrently evolved large bodies, long lifespans, and reduced cancer risk. We also show through comparative genomic analysis that Galapagos giant tortoises have gene duplications related to longevity and tumor suppression. To examine the molecular basis underlying increased body size and lifespan in turtles, we treated cell lines from multiple species, including Galapagos giant tortoises, with drugs that induce different types of cytotoxic stress. Our results indicate that turtle cells, in general, are resistant to oxidative stress related to aging, while Galapagos giant tortoise cells, specifically, are sensitive to endoplasmic reticulum stress, which may give this species an ability to mitigate the effects of cellular stress associated with increased body size and longevity.

## Introduction

Body size and longevity are fundamental life history traits that vary tremendously across vertebrates. Maximum body mass in vertebrates ranges from 0.5 g in the red-backed salamander (*Plethodon cinereus*) (Moore et al., 2001) to 200,000 kg in the blue whale (*Balaenoptera musculus*) (Lockyer, 1976), while maximum lifespan ranges from 8 weeks in the pygmy goby (*Eviota sigillata*) (Depczynski and Bellwood, 2005) to over 400 years in the Greenland shark (*Somniosus microcephalus*) (Nielsen et al., 2016). Life history comparisons also show a strong positive correlation between body size and lifespan across animals, with few exceptions (Healy et al., 2014). There are powerful physiological constraints acting on organisms at the larger, longer-lived end of this spectrum, particularly the accumulation of genetic and cellular damage that comes with having more cells and greater cell turnover (Peto, 2015). The consequences of such long-term genotoxic and cytotoxic stress include genome instability, mitochondrial dysfunction, telomere reduction, and increased cancer risk (López-Otín et al., 2013).

A recurring theme in lifespan and aging regulation is the critical role played by processes that promote cellular protection and maintenance (Kenyon, 2010), including the ability of cells to recycle materials, repair damage, and remove waste. Senescent cells, whose numbers greatly increase with age, exhibit declines in these processes, and are also associated with pro-inflammatory phenotypes that are linked to age-related diseases (Baar et al., 2017; Flatt and Partridge, 2018). At the same time, apoptosis, which is the programmed destruction of unfit or damaged cells, is reduced in older individuals (Salminen et al., 2011). This decline in cell performance in combination with a decreased ability to remove poor performing cells are central to the aging process (López-Otín et al., 2013). Similarly, cancer can arise from cumulative genotoxic and cytotoxic stress, and apoptosis also plays a primary role in cancer resistance by removing potentially cancerous cells (Verfaillie et al., 2013). Thus, if cancer-suppressing mechanisms are similar across species, then larger, longer-lived organisms should be at greater risk of cancer than smaller, shorter-lived ones (Peto, 2015).

The molecular and cellular mechanisms underlying the evolution of large bodies and long lifespans have been explored in mammals such as elephants (Sulak et al., 2016; Vazquez et al., 2018), whales (Seim et al., 2014), bats (Foley et al., 2018; Gorbunova et al., 2020), and naked mole rats (Salmon et al., 2008; Gorbunova et al., 2014), but are less well studied in other vertebrates. Reptiles are an excellent system in which to study the evolution of body size and longevity because diverse lineages have repeatedly evolved large body sizes and long lifespans (Chiari et al., 2018). Turtles, in particular, have lower rates of neoplasia than snakes and lizards (Garner et al., 2004; Sykes and Trupkiewicz, 2006), are especially long-lived, and are “slower aging” than other reptiles (Hoekstra et al., 2020). Most notably, Galapagos giant tortoises (*Chelonoidis niger* species complex; hereafter referred to as *C. niger*) and Aldabra giant tortoises (*Aldabrachelys gigantea*) can live over 150 years (3-5 times longer than their closest relatives) and weigh over 200 kg (50-100 times heavier than their closest relatives) (Caccone et al., 1999; Palkovacs et al., 2002; Poulakakis et al., 2012; Chiari, 2020). Galapagos giant tortoises also appear to have evolved a suite of cellular traits that may contribute to their longevity, such as a slower rate of telomere shortening and extended cellular lifespans compared to mammals (Goldstein, 1974).

Here, we explore the evolution of body size and lifespan in turtles by integrating several approaches (**Figure 1**): (1) phylogenetic comparative analysis of body size, lifespan, and intrinsic cancer risk in turtles; (2) gene duplication analysis of aging and cancer-related genes across available turtle genomes; (3) cell-based assays of apoptosis and necrosis in multiple turtle species varying in body size and lifespan. We show that species with remarkably long lifespans, such as Galapagos giant tortoises, also evolved reduced cancer risk. We also confirm that the Galapagos giant and desert tortoise genomes encode numerous duplicated genes with tumor suppressor and anti-aging functions (Quesada et al., 2019). Our comparative genomic analysis further suggests that cells from large, long-lived species may respond differently to cytotoxic stress, including endoplasmic reticulum (ER) and oxidative stress. The combined genomic and cellular results suggest that at least some turtle lineages evolved large bodies and long lifespans, in part, by increasing the copy number of tumor suppressors and other anti-aging genes and undergoing changes in cellular phenotypes associated with cellular stress.

**Figure 1:**
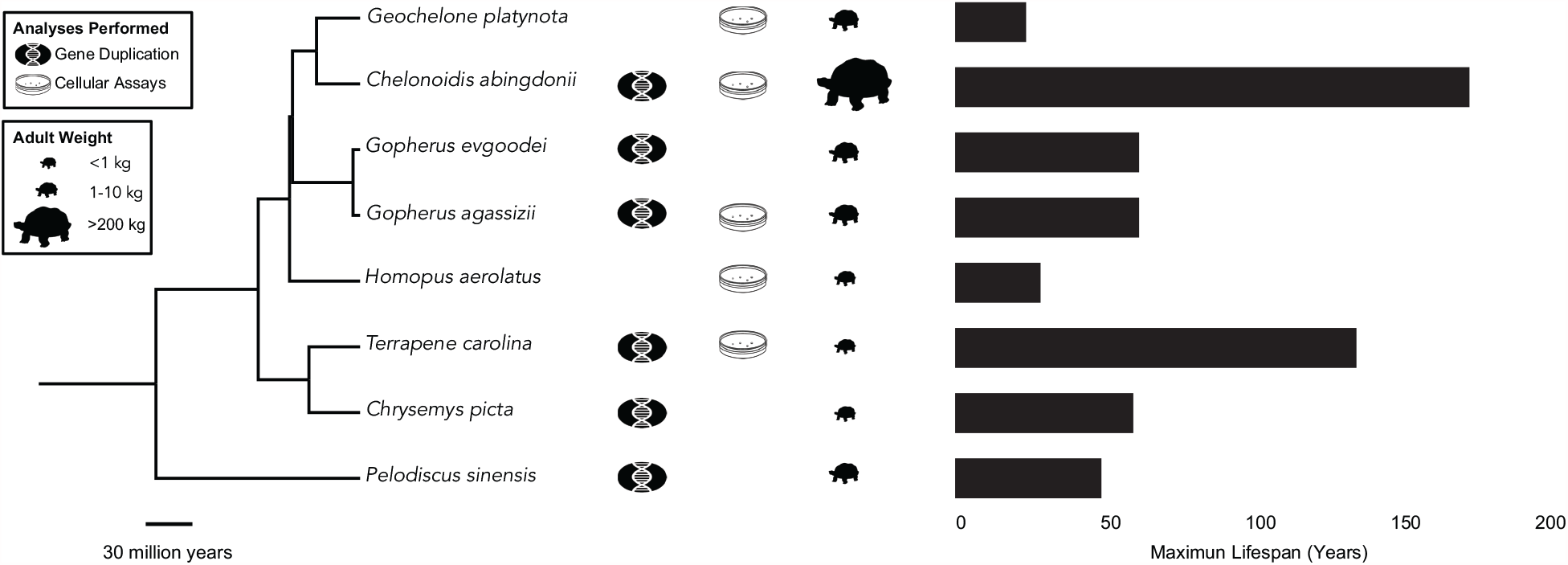
Overview of the study design. Species with genomes utilized for gene duplication analysis as well as turtle species with cells used to measure apoptotic responses to genotoxic and cytotoxic drugs are indicated. Phylogenetic tree was built with TimeTree (Kumar et al., 2017). Turtle size data come from (Ernst and Barbour, 1992; Colston et al., 2020), longevity data are from AnAge.

## Results

### Repeated evolution of large body size in turtles

We found substantial independent accelerations in the rate of body size evolution in several turtle lineages, including a 29x rate increase (386% increase in carapace length) in the stem-lineage of sea turtles (Chelonioidea) and a further 103x rate increase (757% increase in carapace length) in leatherback sea turtles (*Dermochelys coriacea*), a 37x rate increase (35% increase in carapace length) in the stem-lineage of soft-shell turtles (Trionychidae), a 200x rate increase (364% increase in carapace length) in the stem-lineage of narrow-headed softshell turtles (*Chitra chitra* and *Chitra indica*), and a 463x rate increase (364% increase in carapace length) in Cantor’s giant softshell turtle (*Pelochelys cantorii*) (**Figure 2A**).

**Figure 2:**
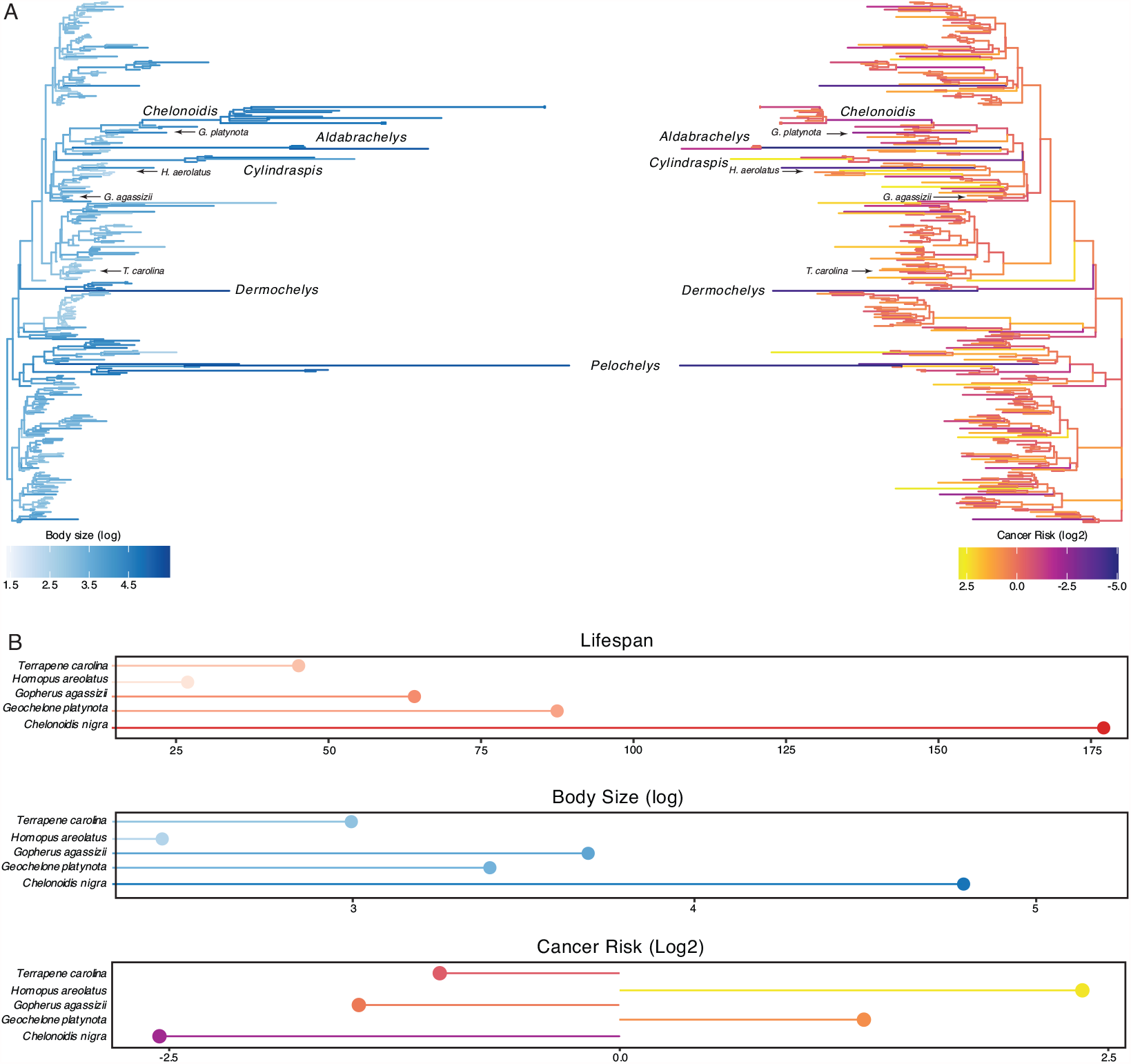
Convergent evolution of large-bodied, cancer resistant turtles. **(A)** Turtle phylogeny with branch lengths and colors scaled by log_2_ change in body size (left) and estimated intrinsic cancer risk (right). Clades and lineages leading to exceptionally large turtles and tortoises are labeled, as are species used in cytotoxic stress assays. **(B)** Lifespan (upper), body size (middle), and estimated intrinsic cancer risk (lower) of species used in stress assays. **Figure 2 – source data 1**. Ancestral reconstructions of testudine body size, lifespan, and intrinsic cancer risk.

Among the more notable groups with increased rates of body size evolution were the “giant” tortoises, including a 81x rate increase (10% increase in carapace length) in the stem-lineage of recently extinct Mascarene giant tortoises (*Cylindraspis* spp.), a 137x rate increase (55% increase in carapace length) in the stem-lineage of Aldabra giant tortoises (*Aldabrachelys* spp.) and a 87x rate increase (383% increase in carapace length) in *Aldabrachelys gigantea*, and a series of rate accelerations in the ancestral lineages of Galapagos giant tortoises, including a 20x rate increase (19% increase in carapace length) in the stem-lineage of *Geochelone* and *Chelonoidis*, a 20x rate increase (27% increase in carapace length) in the stem-lineage of *Chelonoidis*, and a 49x rate increase (81% increase in carapace length) in the stem-lineage of *C. niger*. These data indicate that gigantism evolved independently in multiple lineages of turtles, and step-wise in the evolution of giant tortoises with several rate accelerations in lineages ancestral to *C. niger*.

### Reduction of intrinsic cancer risk in turtles

In order to account for a relatively constant prevalence of cancer across species (Dorn et al., 1968; Abegglen et al., 2015; Boddy et al., 2020), intrinsic cancer risk must coevolve with changes in body size and lifespan across species. For example, a 100-year retrospective study of neoplasia in zoo reptiles identified only six neoplasms in 490 turtle necropsies, which ranged in size from the West African mud turtle (*Pelusios castaneus*, carapace length ca. 25–28 cm) to the spiny softshell turtle (*Apalone spinifer spinifer*, carapace length ca. 54 cm) (Sykes and Trupkiewicz, 2006). As expected, relative intrinsic cancer risk (RICR) in turtles also varies with changes in body size and lifespan (**Figure 2A**). We estimated a 73-log_2_ decrease RICR in the stem-lineage of sea turtles (Chelonioidea), a 129-log_2_ decrease RICR in leatherback sea turtles (*Dermochelys coriacea*), a 14-log_2_ decrease RICR in the stem-lineage of soft-shell turtles (Trionychidae), a 34-log_2_ decrease RICR in the stem-lineage of narrow-headed softshell turtles (*Chitra chitra* and *Chitra indica*), and a 140-log_2_ decrease RICR in the stem-lineage of Cantor’s giant softshell turtle (*Pelochelys cantorii*).

Among the “giant” tortoises, we estimated a 97-log_2_ decrease RICR in the stem-lineage of Mascarene giant tortoises (*Cylindraspis* spp.), a 154-log_2_ decrease RICR in the stem-lineage of Aldabra giant tortoises (*Aldabrachelys* spp.) and a 50-log_2_ decrease RICR in *Aldabrachelys gigantea*. In the lineages ancestral to Galapagos giant tortoises, we estimated a 67-log_2_ decrease RICR in the stem-lineage of *C. niger* a 27-log_2_ decrease RICR in the stem-lineage of the *Chelonoidis*, and a 19-log_2_ decrease RICR in the stem-lineage of *Geochelone* and *Chelonoidis*. Thus, turtles coevolved large bodies and reduced intrinsic cancer risk, including step-wise reductions in the lineages ancestral to *C. niger*.

### Identification of tumor suppressor and anti-aging gene duplications in turtle genomes

Previous studies have shown that large-bodied cancer resistant species such as elephants (Sulak et al., 2016; Vazquez and Lynch, 2021) and whales (Keane et al., 2015) evolved an increased number of tumor suppressors, suggesting that the same may be possible in giant, long-lived turtles. A previous study of Galapagos giant tortoises, for example, identified several gene duplications in pathways that might be related to body size evolution and reduced cancer risk (Quesada et al., 2019). Therefore, we reanalyzed the Galapagos giant tortoise genome and other turtle genomes to identify gene duplications and used maximum likelihood-based ancestral state reconstruction to determine lineages in which genes were duplicated. We identified ∼86 duplications in the stem-lineage of Testudines, 245 in the stem-lineage of Pleurodira, 33 in the stem-lineage of tortoises (AncTortoise), 259 in *C. abingdonii*, 201 in *G. agassizii*, 273 in *T. carolina*, 315 in *C. picta*, and 270 in *P. sinensis* (**Figure 3A**).

**Figure 3:**
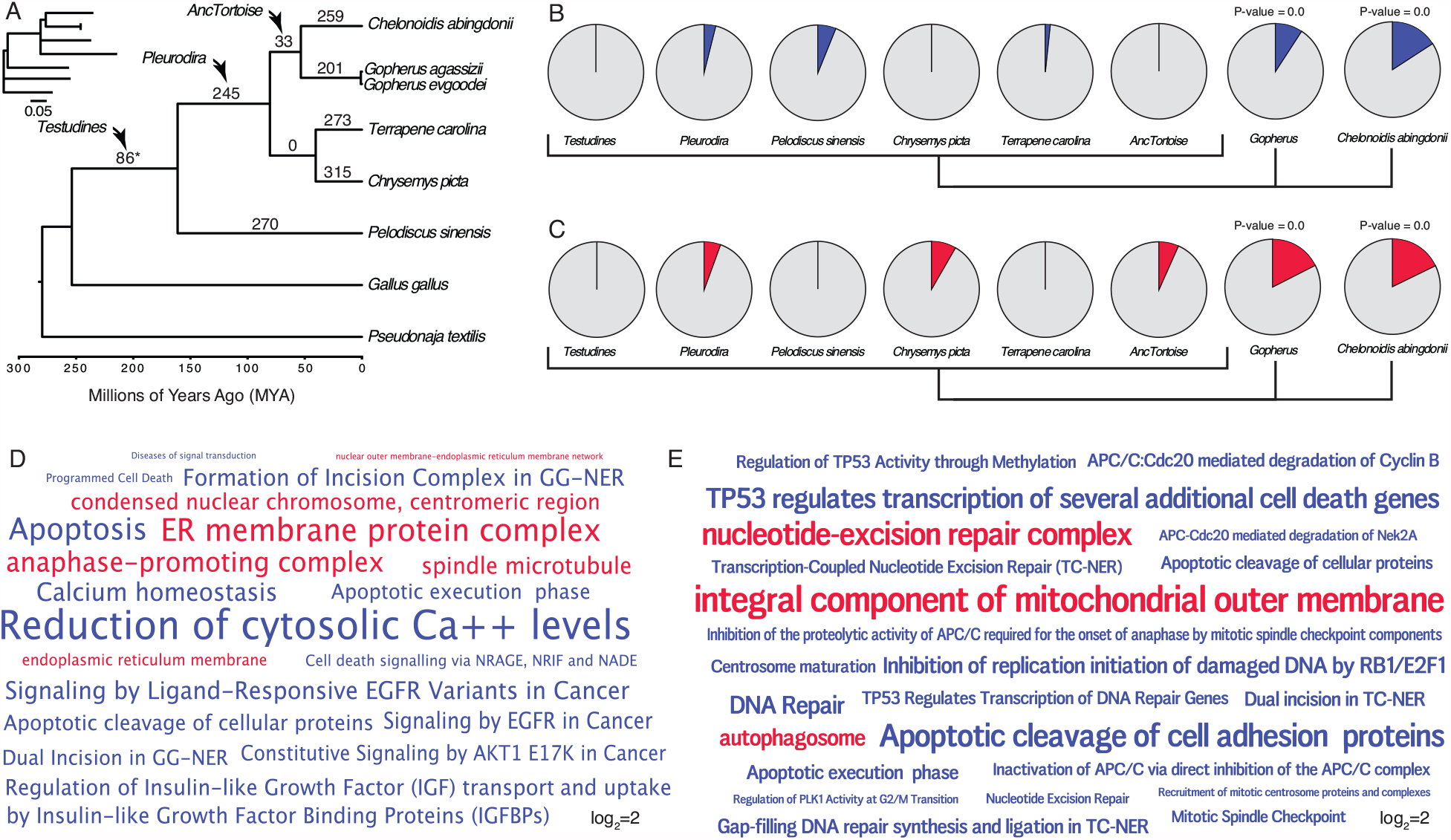
Gene duplicates in Galapagos and desert tortoises are enriched in tumor suppressor and anti-aging functions. **(A)** Turtle phylogeny indicating the number of genes duplicated in each lineage, inferred by maximum likelihood. Inset, phylogeny with branch lengths proportional to gene duplication rate. The asterisk (*) denotes a node with gene duplications reconstructed with lower support than other nodes and non-significant (BPP=0.541). **(B)** Pie charts indicating the proportion of enriched Reactome pathways in each lineage related to cancer biology and aging (blue slices). Gene duplicates in Galapagos giant and desert tortoises are significantly more enriched in these terms than other lineages (two-sided permutation t-test is 0.00). **(C)** Pie charts indicating the proportion of enriched GO cellular component terms related to cancer biology, DNA damage repair, programmed cell death, and the endoplasmic reticulum (red slices). Gene duplicates in Galapagos giant and desert tortoises are significantly more enriched in these terms than other lineages (two-sided permutation t-test is 0.00). **(D)** Wordcloud of the Reactome (blue) pathways and GO cellular component terms (red) enriched exclusively in Galapagos giant tortoises. Only pathway and GO terms enriched with P<=0.05 are shown are scaled according to Log2 fold-enrichment (see inset scale). **(E)** Wordcloud of the Reactome (blue) pathways and GO cellular component terms (red) enriched in desert tortoises. Only pathway and GO terms enriched with P<=0.05 are shown are scaled according to Log2 fold-enrichment (see inset scale). **Figure 3 – source data 1**. Ancestral reconstruction of copy number changes. **Figure 3 – source data 2**. Gene duplications and pathway enrichments for each lineage.

Consistent with previous studies which observed duplication of tumor suppressor and other anti-aging genes in large, long-lived species, we found that 12% of the pathways enriched among Galapagos giant tortoises were related to cancer and aging biology, while only 0-6% of the pathways that were enriched among gene duplications in other lineages were related to cancer and aging biology (**Figure 3B**). Next, we identified GO cellular component terms that were enriched among gene duplications in each lineage. While gene duplications in some lineages were enriched in GO terms related to cancer biology and aging, significantly more ontology terms in Galapagos giant tortoises were related to cancer and aging biology (**Figure 3C**). Enriched pathway and ontology terms in Galapagos (**Figure 3D**) tortoises included “Apoptosis”, “Programmed Cell Death”, “Cell death signalling via NRAGE, NRIF and NADE”, “Dual Incision in GG-NER’’ and “Formation of Incision Complex in GG-NER”, and “Reduction of cytosolic Ca++ levels”. We also observed that “Regulation of Insulin-like Growth Factor (IGF) transport and uptake by Insulin-like Growth Factor Binding Proteins (IGFBPs)” was an enriched pathway term among Galapagos giant tortoise gene duplications, which may be related to the regulation of body size. Among the GO cellular component terms exclusively enriched among Galapagos giant tortoise gene duplications were “ER membrane protein complex”, “endoplasmic reticulum membrane” and “nuclear outer membrane-endoplasmic reticulum membrane network”, and “anaphase-promoting complex”.

Desert tortoise specific gene duplications were enriched in pathways related to cancer biology and aging, particularly compared to other turtles. For example, 9.4% of the pathways enriched among desert tortoise duplicates were related to cancer and aging biology, significantly more than other turtle lineages but less than Galapagos giant tortoises (**Figure 3B**). Similarly, significantly more GO terms in desert tortoises were related to cancer and aging biology (**Figure 3C**). Enriched pathway and ontology terms in desert tortoises (**Figure 3E**) were related to DNA damage and repair including “Formation of TC-NER Pre-Incision Complex”, “Gap-filling DNA repair synthesis and ligation in TC-NER”, “Dual incision in TC-NER”, “Apoptotic cleavage of cell adhesion proteins”, and “Transcription-Coupled Nucleotide Excision Repair (TC-NER)”. Enriched in GO terms included “integral component of mitochondrial outer membrane”, “nucleotide-excision repair complex”, and “autophagosome”. These data suggest that desert tortoises have evolved gene duplications that may also contribute to cancer resistance and the evolution of longevity.

### Turtle cells have unique responses to genotoxic and cytotoxic stress

Our observation that gene duplications in the Galapagos and desert tortoise genomes are enriched in pathways and GO terms related to the biology of aging, apoptosis, cell cycle regulation, DNA damage repair, and mitochondrial oxidative DNA damage protection (**Figure 3D**), suggests that cells from these species may have different cellular responses to genotoxic and cytotoxic stress than cells from other turtles. To test this hypothesis, we treated primary fibroblasts from *C. niger, G. platynota, G. agassizii, H. aerolatus*, and *T. carolina* (**Figure 2B**) with drugs to induce different types of stress including: 1) tunicamycin, which induces endoplasmic reticulum (ER) stress and the unfolded protein response (UPR) through an accumulation of unfolded and misfolded proteins (Banerjee et al., 2011; Guha et al., 2017); 2) etoposide, which forms a ternary complex with DNA and topoisomerase II and prevents re-ligation of replicating DNA strands leading to single- and double-stranded DNA breaks (Wozniak and Ross, 1983); and 3) paraquat, which causes oxidative stress through the production of reactive oxygen species and S-phase cell cycle arrest (Salmon et al., 2008). We quantified the kinetics of cell death using the RealTime-Glo™ Annexin V Apoptosis and Necrosis assay (RTG) every 30 minutes for 48 hours.

We found that tunicamycin induced a dose dependent increase in apoptosis in cells from most species (**Figure 4A**). However, at 24 hrs and 25 uM, *C. niger* cells had an apoptotic response that was at least double that of other species while *G. platynota* cells were insensitive to tunicamycin (**Figure 4A**). In contrast etoposide did not induce apoptosis or necrosis in cells from most species but did induce a strong apoptotic response in *G. agassizii* cells (**Figure 4B**). Turtle cells were also variably sensitive to paraquat, but cells from all species induced apoptosis in response to paraquat treatment (**Figure 4C**). Thus, *C. niger* cells are more sensitive to ER and UPR stress than other species, *G. agassizii* cells are more sensitive to DNA damage induced by etoposide than other species, and cells from all species were sensitive to oxidative stress induced by paraquat.

**Figure 4:**
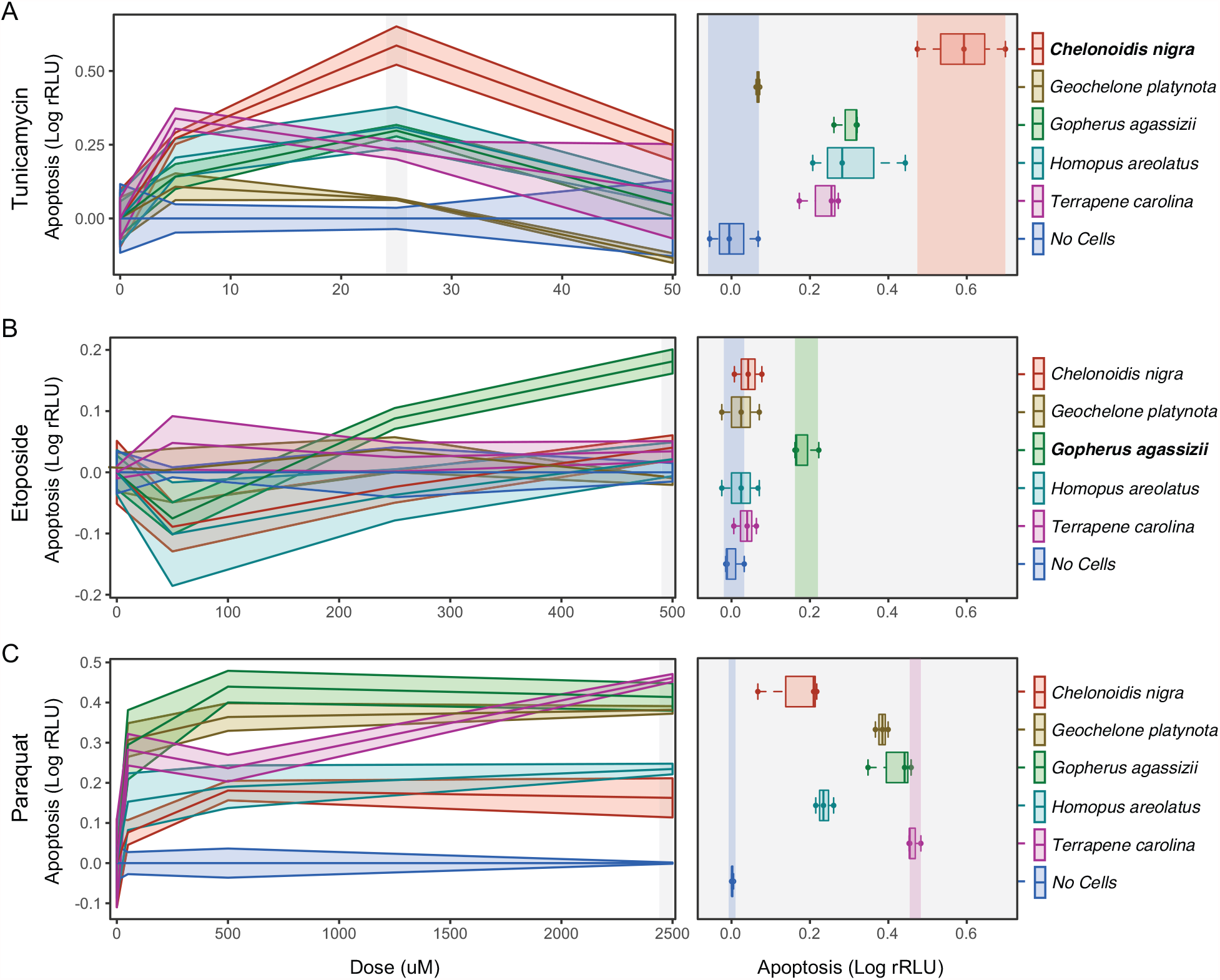
Cells from Galapagos giant and desert tortoises have unique stress responses. **(A)** Left, dose response curves for tunicamycin, which induces endoplasmic reticulum stress and the unfolded protein response (24 hours post-treatment). Right, boxplots showing differences between species 24 hours after treatment with 25uM tunicamycin; statistical tests are relative to *C. niger*, which had the strongest apoptotic response. The unpaired mean difference between *C. niger* and *G. platynota* is -0.521 [95.0%CI -0.631, -0.408]. The *P* value of the two-sided permutation t-test is 0.0. The unpaired mean difference between *C. niger* and *G. agassizii is* -0.288 [95.0%CI -0.399, -0.193]. The *P* value of the two-sided permutation t-test is 0.0. The unpaired mean difference between *C. niger* and *H. areolatus is* -0.278 [95.0%CI -0.431, -0.124]. The *P* value of the two-sided permutation t-test is 0.0. The unpaired mean difference between *C. niger* and *T. carolina* is -0.355 [95.0%CI -0.465, -0.246]. The *P* value of the two-sided permutation t-test is 0.0. n=3. **(B)** Left, dose response curves for etoposide, which induces DNA strand breaks (24 hours post-treatment). Right, boxplots showing differences between species 24 hours after treatment with 500uM etoposide; statistical tests are relative to *G. agassizii*, which had the strongest apoptotic response. The unpaired mean difference between *G. agassizii* and *C. niger* is -0.141 [95.0%CI -0.195, -0.106]. The *P* value of the two-sided permutation t-test is 0.0.The unpaired mean difference between *G. agassizii* and *G. platynota* is -0.16 [95.0%CI -0.227, -0.112]. The *P* value of the two-sided permutation t-test is 0.0.The unpaired mean difference between *G. agassizii* and *H. areolatus* is -0.16 [95.0%CI -0.227, -0.112]. The *P* value of the two-sided permutation t-test is 0.0. The unpaired mean difference between *G. agassizii* and *T. carolina* is -0.147 [95.0%CI - 0.198, -0.115]. The *P* value of the two-sided permutation t-test is 0.0. n=3. **(C)** Left, dose response curves for paraquat, which induces oxidative stress (24 hours post-treatment). Right, boxplots showing differences between species 24 hours after treatment with 2500uM paraquat; statistical tests are relative to *C. niger*, which had the weakest apoptotic response. The unpaired mean difference between *C. niger* and *G. platynota* is 0.24 [95.0%CI 0.176, 0.317]. The P value of the two-sided permutation t-test is 0.0. The unpaired mean difference between *C. niger* and *G. agassizii* is 0.272 [95.0%CI 0.188, 0.356]. The P value of the two-sided permutation t-test is 0.0. The unpaired mean difference between *C. niger* and *H. areolatus* is 0.0927 [95.0%CI 0.0337, 0.169]. The P value of the two-sided permutation t-test is 0.0. The unpaired mean difference between *C. niger* and *T. carolina* is 0.32 [95.0%CI 0.262, 0.396]. The P value of the two-sided permutation t-test is 0.0. n=3 **Figure 4 – figure supplement 1**. Tunicamycin RealTime-Glo time course. **Figure 4 – figure supplement 2**. Etoposide RealTime-Glo time course. **Figure 4 – figure supplement 3**. Paraquat RealTime-Glo time course. **Figure 4 – source data 1**. RealTime-Glo datafiles.

## Discussion

Reptiles in general, and turtles specifically, are an excellent system for studying the mechanisms underlying variation in body size, lifespan, and cancer resistance (Chiari et al., 2018; Hoekstra et al., 2020). Adult body size among turtle species differs by three orders of magnitude, ranging from the speckled dwarf tortoise (*Chersobius signatus*; 100 g) to Aldabra giant tortoises (*A. gigantea*; >300 kg), while longevity across species varies from tens of years to >150 years in giant tortoises. Yet, even the smallest turtles live relatively long (up to 30 years) compared to most other vertebrates. Turtles also have lower estimated cancer rates (∼1.2%) compared to mammals (∼12.5%), suggesting that they have evolved the means to delay aging and reduce cancer susceptibility (Garner et al., 2004; Sykes and Trupkiewicz, 2006; Abegglen et al., 2015; Boddy et al., 2020). While increased lifespan and cancer prevalence in turtles could, in part, be due to reduced damage resulting from their lower metabolic rates, previous genomic and cellular data suggests that there may also be molecular differences that enable extremes in body size, longevity, and cancer resistance (Goldstein, 1974; Quesada et al., 2019).

Here, we found that body size rapidly increased in multiple turtle lineages independently, including a dramatic increase in both body size and the rate of body size evolution in Galapagos giant tortoises. When body size, longevity, and intrinsic cancer rates are analyzed together within an explicit phylogenetic context, large-bodied lineages evolved reduced intrinsic cancer risk. Among the most dramatic decreases in cancer risk among turtles was in the stem-lineage of Galapagos giant tortoises, suggesting this lineage evolved genetic and cellular mechanisms that reduce cancer risk. Consistent with the evolution of reduced intrinsic cancer risk in this lineage, we found that gene duplications in the Galapagos lineage are enriched in ER stress associated pathways is similar to a previous analysis by Quesada et al. (2019), who found that genes under positive selection in Galapagos giant tortoises are also enriched for ER function and stress pathways. ER stress dysregulates protein homeostasis, which is one of the hallmarks of aging and reduced animal lifespan (López-Otín et al., 2013; Morimoto and Cuervo, 2014; Kaushik and Cuervo, 2015). The impairment of protein production, folding, and degradation associated with ER stress can impact many cellular processes and lead to the buildup of toxic protein aggregates that are linked to many age-related diseases and cancers (Vilchez et al., 2014).

Our data suggest a mechanistic connection between genes related to the ER, enhanced responses to ER stress, and the UPR in Galapagos giant tortoise cells. Indeed, we found that tunicamycin, which induces ER stress by activating the UPR and ultimately inducing apoptosis (Nami et al., 2016), causes an immediate and pronounced apoptotic response in Galapagos giant tortoise cells, while cells from other species were much slower to respond. These results indicate that Galapagos giant tortoise cells have an extremely sensitive ER stress response. In contrast, Galapagos giant tortoise cells did not exhibit a heightened apoptotic response to paraquat and etoposide, which induce oxidative stress, DNA damage, and apoptosis independently of the ER stress/UPR signaling pathway (Mizumoto et al., 1994; Suntres, 2002). These data suggest that changes in the ER stress and UPR signaling pathways may contribute to the evolution of long lifespans, large bodies, and augmented cancer resistance in Galapagos giant tortoises.

Unexpectedly, we also found that cells from most species were unresponsive to etoposide treatment, even at the highest dose and longest exposure time. The sole exception was *G. agassizii* cells, which were similar to other species in their response to tunicamycin and paraquat, but markedly more sensitive to etoposide treatment. These data suggest that turtle cells can either export intra-cellular etoposide before it can induce DNA damage, are generally insensitive to DNA damage induced by etoposide, or can rapidly repair etoposide induced DNA damage. Regardless of these, and potentially other mechanisms that alter sensitivity to etoposide, *G. agassizii* cells respond differently than cells from the other species tested. The functional and organismal consequences of this altered sensitivity are unclear, but is similar to elephant cells which have evolved to induce apoptosis at relatively low levels of DNA damage(Sulak et al., 2016; Vazquez et al., 2018; Vazquez and Lynch, 2021). This reduced threshold for inducing apoptosis in response to cellular stress may clear cells that have been exposed to the kinds of stresses that eventually lead to cancer before transformation into cancer cells.

Although medical literature has often portrayed high levels of apoptosis as a maladaptive response to cellular stress contributing to aging (Szegezdi et al., 2006; Chadwick and Lajoie, 2019), previous work in elephants showed that heightened apoptotic responses to genetic and cellular damage can actually be an adaptive and beneficial response linked to increased body size and longevity (Abegglen et al., 2015; Sulak et al., 2016; Vazquez et al., 2018). This is because rapid and effective clearance of damaged or injured cells can help maintain tissue integrity (Baar et al., 2017; de Keizer, 2017), especially in organisms with many cells and large amounts of cell turnover. Thus, our results are compatible with ER stress as a potential factor in the evolution of large, long-lived turtles.

### Caveats and limitations

There are several limitations to our findings that could be the subject of future research. While all cellular assays were performed on a common cell type, fibroblasts, the tissue origin of these cells differed among species. We note, however, that for Galapagos giant tortoises, tunicamycin response was not confounded with the site of fibroblast origin. We were also unable to obtain the age of source animals, which could affect results since apoptosis can decline during senescence (Salminen et al., 2011). Finally, the taxon sampling can be expanded to include cells from closely related species with large differences in body size or lifespan. This would enable better resolution in isolating the evolutionary origins of enhanced responses to genetic and cellular stress, for example, by including the closest living relative of Galapagos giant tortoises (Chaco tortoises; *Chelonoidis chilensis*). We relied on primary cell lines that were currently available in frozen zoos and commercial biobanks; but future cellular work could attempt to sample fresh cells from the same tissue type and life stage across species, which is logistically challenging.

### Conclusions

While numerous studies have found comparative genomic signatures associated with the evolution of body size, longevity, and cancer resistance (Keane et al., 2015; Herrera-Álvarez et al., 2018; Babarinde and Saitou, 2020), there have been few attempts to validate these findings experimentally at the cellular level (Jimenez et al., 2018). Furthermore, most previous cellular studies on these subjects focus almost exclusively on placental mammals. The work presented here utilizes turtle cell lines, which is much more feasible than studying body size and aging phenotypes in these long-lived animals. Our most salient finding is that Galapagos giant tortoises are much more sensitive at inducing apoptosis in response to ER stress compared to other turtle species, and also have genomic and phylogenetic signatures of rapid evolutionary increases in body size, lifespan, and cancer resistance. We also found more generally that all turtle cell lines were resistant to oxidative stress induced by paraquat. This supports previous oxidative stress studies in turtles, and indicates that turles, in general, may be a promising model system in which to study resistance to stress from long lifespans (Lutz et al., 2003). However, while the resistance of turtle cells to oxidative stress may contribute to the generally long lifespans of turtles, the sensitivity of Galapagos giant tortoise cells to ER stress may increase their resistance to oncogenic transformation thus promoting healthy aging and larger body sizes.

## Methods

### Intrinsic cancer risk estimation

The dramatic increase in body size and lifespan in some turtle lineages, and the relatively constant rate of cancer across species of diverse body sizes and lifespan (Leroi et al., 2003), would predict an increase in cancer risk concurrent with an increase in body size or lifespan. In order to identify lineages with exceptional changes in body size (with total carapace length used here as a proxy for body size), longevity, or intrinsic cancer risk, we jointly estimated these parameters across turtles and reconstructed ancestral states within a phylogenetic framework. Using phylogenetic and body size data from Colston et al. (2020) and longevity data from AnAge (Magalhães and Costa, 2009), and following Peto’s model of cancer risk (2015; Vazquez and Lynch, 2021), we estimated the intrinsic cancer risk (*K*) as the product of risk associated with body size and lifespan (lifespan^6^ x body size). In order to determine (*K*) across species and at ancestral nodes, we first estimated body size at each node. We used a generalization of the Brownian motion model that relaxes assumptions of neutrality and gradualism by considering increments to evolving characters to be drawn from a heavy-tailed stable distribution (the “stable model”) implemented in StableTraits (Elliot and Mooers, 2014). The stable model allows for large jumps in traits and has previously been shown to out-perform other models of body size evolution, including standard Brownian motion models, Ornstein– Uhlenbeck models, early burst maximum likelihood models, and heterogeneous multi-rate models (Prang, 2019). We used Phylogenetic Generalized Least-Square Regression (PGLS) (Grafen and Hamilton, 1989; Martins and Hansen, 1997; Pagel, 1997) using a Brownian covariance matrix as implemented in the R package *ape* (Paradis and Schliep, 2019) to infer ancestral lifespans across turtles using our estimates for body size (Colston et al., 2020) and reported maximum lifespans for each species (Magalhães and Costa, 2009). Fold-change in cancer susceptibility was estimated between all ancestral (K1) and descendant (K2) nodes. The fold change in cancer risk between a node and its ancestor was then defined as K2/K1 (Figure 2 – source data 1).

### Identification of duplicated genes and reconstruction of ancestral copy numbers

Following Caulin (2015), we identified duplicated genes in the genomes of the Pinta Island Galapagos giant tortoise (*Chelonoidis abingdonii*; ASM359739v1), Goode’s thornscrub tortoise (*Gopherus evgoodei*; rGopEvg1_v1.p) and Agassiz’s desert tortoise (*Gopherus agassizii*; ASM289641v1), Chinese softshell turtle (*Pelodiscus sinensis*; PelSin_1.0), painted turtle (*Chrysemys picta bellii*; Chrysemys_picta_bellii-3.0.3), three-toed box turtle (*Terrapene carolina triunguis*; T_m_triunguis-2.0), chicken (Gallus gallus; GRCg6a), and eastern brown snake (*Pseudonaja textilis*; EBS10Xv2-PRI) using the Ensembl (Genes 103) BioMart web-based tool to extract same-species paralogies (within_species_paralog) from each genome; same-species paralogies are identified by Ensembl Compara (Howe et al., 2021) using gene tree species tree reconciliation. Genes that were classified as pseudogenes on Ensembl were not included. Duplicate genes identified in Goode’s thornscrub tortoise (*G. evgoodei*; rGopEvg1_v1.p) were manually verified in Agassiz’s desert tortoise (*G. agassizii*; ASM289641v1) using reciprocal best BLAT.

We used maximum likelihood-based ancestral state reconstruction to determine when in the evolution of turtles each gene was duplicated. We encoded the copy number of each putatively functional gene for each species as a discrete trait, with state 0 for one gene copy and state 1 for two or more copies. We used IQ-TREE to select the best-fitting model of character evolution (Minh et al., 2020; Hoang et al., 2018; Kalyaanamoorthy et al., 2017; Wang et al., 2018; Schrempf et al., 2019), which was inferred to be a general time reversible model for morphological data (GTR2) with character state frequency optimized (FO) by maximum-likelihood from the data. Next, we inferred gene duplication events with the empirical Bayesian ancestral state reconstruction (ASR) method implemented in IQ-TREE (Minh et al., 2020; Hoang et al., 2018; Kalyaanamoorthy et al., 2017; Wang et al., 2018; Schrempf et al., 2019), the best fitting model of character evolution (GTR2+FO) (Soubrier et al., 2012; Yang et al., 1995), and the unrooted species tree for turtles (**Figure 3A**). We considered ancestral state reconstructions to be reliable if they had Bayesian Posterior Probability (BPP) ≥ observed state frequency from the alignment; less reliable reconstructions were excluded from further analyses.

### Gene duplication pathway enrichment analysis

To determine if gene duplications were enriched in particular biological pathways, we used Enrichr (Chen et al., 2013, p 5; Kuleshov et al., 2016) to perform Over-Representation Analysis (ORA) of the Reactome database (Jassal et al., 2020). Furthermore, we used the Panther GO enrichment analysis tool (Mi et al., 2021) to perform ORA on GO cellular component terms (Ashburner et al., 2000; The Gene Ontology Consortium, 2021). Gene duplicates in each lineage were used as the foreground gene set, and the initial query set was used as the background gene set. Enrichr uses a hypergeometric test for statistical significance of pathway over-representation, while the Panther GO enrichment analysis tool uses a binomial test for statistical significance of GO cellular component term over-representation.

### Cell culture

Experimental cellular phenotypes were generated to compare results from the body size, longevity, and intrinsic cancer risk analysis and the gene duplication and enrichment analysis. Apoptosis was chosen as the major cellular endpoint of interest because of its central role in aging and cancer through the removal of damaged or cancerous cells (Salminen et al., 2011; Verfaillie et al., 2013).

We cultured cell lines from four tortoise taxa (*Chelonoidis niger, Geochelone platynota, Gopherus agassizii, Homopus aerolatus*) obtained from the San Diego Frozen Zoo and one turtle species (*Terrapene carolina*) obtained from the American Type Culture Collection **(Table 1)**. All cells were primary fibroblasts derived from either the heart (*T. carolina*), trachea (*G. agassizii, G. platynota, C. niger*), or eye (*H. aerolatus*). Turtle cells have previously been shown to grow within a range of 23-30 °C (Clark and Karzon, 1967; Clark et al., 1970; Goldstein, 1974). Therefore, cells were incubated at 25 °C, with 5% CO_2_. Cells were cultured in Minimum Essential Medium (MEM; Gibco) with 10% fetal bovine serum (Gibco) and 1% penicillin-streptomycin antibiotic (Gibco) in standard T75 flasks (Thermo Fisher Scientific). Media was changed every three days. Cells were passaged before reaching 90% confluency, approximately every 7 to 9 days. For passaging, cell plates were rinsed with one volume of 37 °C DPBS (Gibco) and cells detached with 0.25% Trypsin-EDTA (Gibco). We note that cells detached quickly with the assistance of gentle tapping of plates and without incubation. The cell suspension was transferred to a 15-mL conical tube (Thermo Fisher Scientific) with an equal volume of complete media to stop trypsinization. Cells were then centrifuged at 500 x g for 5 minutes, and then the pellet was resuspended in 1 mL of complete media. Cell viability was determined using a TC10 Automated Cell Counter (Bio-Rad Laboratories) and was >75% for all cell lines throughout the experiment.

**Table 1.**
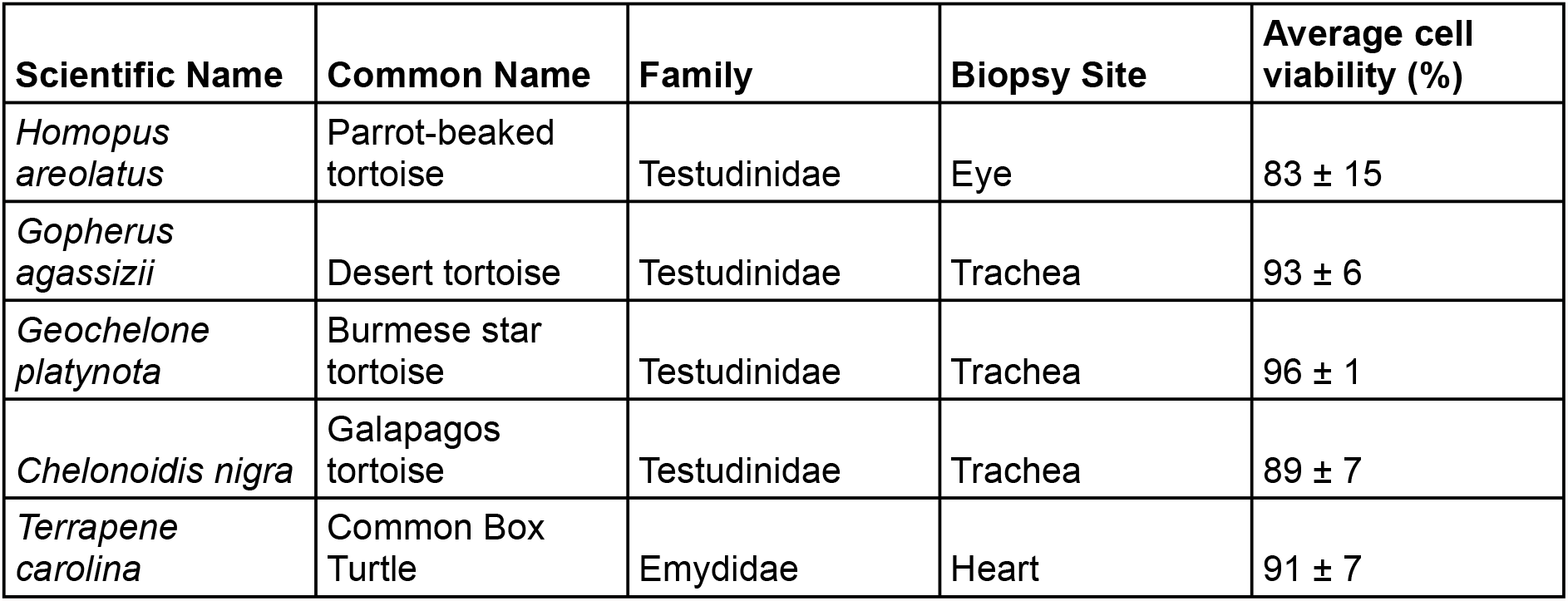
Turtle experimental cell lines. Biopsy site is the location from which primary fibroblast cells were derived. Average *in vitro* cell viability percent is over 12-14 passages depending on cell line. Cell viability was calculated as the ratio of live cells to total cells at each passage with standard deviation.

We selected various cytotoxic drugs to induce different types of cellular stress, including etoposide (Cayman Chemical Company), which induces single-stranded and double stranded DNA breaks (Wozniak and Ross, 1983), paraquat (Sigma-Aldrich), which induces oxidative stress through production of reactive oxygen species and s-phase cell cycle arrest (Salmon et al., 2008), and tunicamycin (Cayman Chemical Company), which induces ER stress and the unfolded protein response (UPR) by causing an accumulation of unfolded and misfolded proteins (Banerjee et al., 2011; Guha et al., 2017).

### Kinetic measurements of cell death

The RealTime-Glo™ Annexin V Apoptosis and Necrosis assay (RTG) visualizes the kinetics of apoptosis over a given period of time and differentiates secondary necrosis occurring during late apoptosis from necrosis caused by other cytotoxic events (Landreman et al., 2019). RTG assays were performed for each species by seeding 5,000 cells per well into an opaque bottomed 96-well plate with three replicates per treatment per species and one empty column with no cells (background control). Cells were left to adhere for 24 hours, after which seeding media was aspirated off and serial dilutions of Tunicamycin (0 μM, 10 μM, 50 μM, 100 μM), Etoposide (0 μM, 100 μM, 500 μM, 1000 μM), or Paraquat (0 μM, 100 μM, 1000 μM, 5000 μM) were applied. All drug treatments were made using Fluorobrite DMEM media. The “0 μM” control treatments consisted of the vehicle used (DMSO or PBS) at the concentration matching the highest drug concentration, while the background control (“NoCell”), consisted of the drug treatment and assay reagents with no cells. As per the RTG product protocol, the 500-fold dilution of reagents in Fluorobrite DMEM media was added to wells immediately after drug treatments were applied (effectively reducing the initial drug dilutions in half). Readings were then taken every 30 minutes for 48 hours using a GloMax Luminometer (Promega Corporation).

### Statistical analysis

All statistical analyses reported in this paper are estimation statistics, including effect sizes, 95% confidence intervals (CIs) of the effect size, and *P* values. Effect sizes and 95% CIs are reported as: effect size [CI width lower bound; upper bound] with 5000 bootstrap samples; the confidence interval is bias-corrected and accelerated. *P* values reported are the likelihood of observing the effect sizes if the null hypothesis of zero difference is true. For each permutation *P* value, 5000 reshuffles of the control and test labels were performed (Ho et al., 2019). All statistical analyses were performed in RStudio (RStudio Team, 2020).

## Acknowledgements

We thank the San Diego Frozen Zoo for providing cell lines for four of the tortoise species in this project. The University of Chicago kindly hosted SG and SEB as part of NSF’s EPSCoR Program.

