## Supplementary material for "Concurrent evolution of anti-aging gene duplications and cellular phenotypes in long-lived turtles": Source data files and figure supplements: parsimony.py

```
#!/usr/bin/python

# This script will read in files containing presence and expression data and produce
# a NEXUS file for parsimony analysis based on these data

# Reference list is the first entry
taxon_list = ['Human','Anole','Snake','Gallus','Chelonoidisabingdonii-within_species_paralog','Chrysemyspicta-within_species_paralog','Gopherusevgoodei-within_species_paralog','Pelodiscussinensis-within_species_paralog','Terrapenecarolinatriunguis-within_species_paralog']

# Create list of lists with all non-human taxa annotated genes
annotated_list = []
for BA in taxon_list:
	temp_list = []
	for line in open('/Users/Vinny/Desktop/Turtledata/Duplicates/within_species_paralog/'+str(BA)+'.annotations.txt','r'):
		AB = line.rstrip()
		AC = AB.upper()
		temp_list.append(AC)
	annotated_list.append(temp_list)
	if BA == 'Human':
		Human_Reference_List = temp_list
	
# Create list of lists with all non-human taxa expressed genes
expressed_list = []
for BA in taxon_list:
	temp_list = []
	for line in open('/Users/Vinny/Desktop/Turtledata/Duplicates/within_species_paralog/'+str(BA)+'.expressed.txt','r'):
		AB = line.rstrip()
		AC = AB.upper()
		temp_list.append(AC)
	expressed_list.append(temp_list)

# Compare lists to sets to determine presence/absence
dataset = []
j = 0
for CA in taxon_list:
	temp_list = []
	annotated_set = set(annotated_list[j])
	expressed_set = set(expressed_list[j])
	for CB in annotated_list[0]:		
		if (CB in annotated_set and CB in expressed_set): temp_list.append("1")
		elif CB in annotated_set: temp_list.append("0")
		else: temp_list.append("?")
	j = j + 1
	dataset.append(temp_list)


# Write out NEXUS file to 'outfile' containing expression data
outfile = open("/Users/Vinny/Desktop/Turtledata/Duplicates/within_species_paralog/Matrix.nex",'w')
outfile.write("#NEXUS\n\nBegin DATA;\n\tDimensions ntax=13 nchar=19656;\n\tFormat datatype=STANDARD gap=- missing=?;\n\tMatrix\n\n")

i = 0
for DA in taxon_list:

	if len(str(DA)) <= 3: outfile.write(DA+"\t\t\t")
	elif len(str(DA)) > 7: outfile.write(DA+"\t")
	else: outfile.write(DA+"\t\t")
	
	for DB in dataset[i]:
		outfile.write(str(DB))
	outfile.write("\n")
	i = i + 1
outfile.write("\n;\nEnd;\n")
outfile.close()
```
