## Supplementary figures and images for "Concurrent evolution of anti-aging gene duplications and cellular phenotypes in long-lived turtles"

### Constraint.tree.pdf

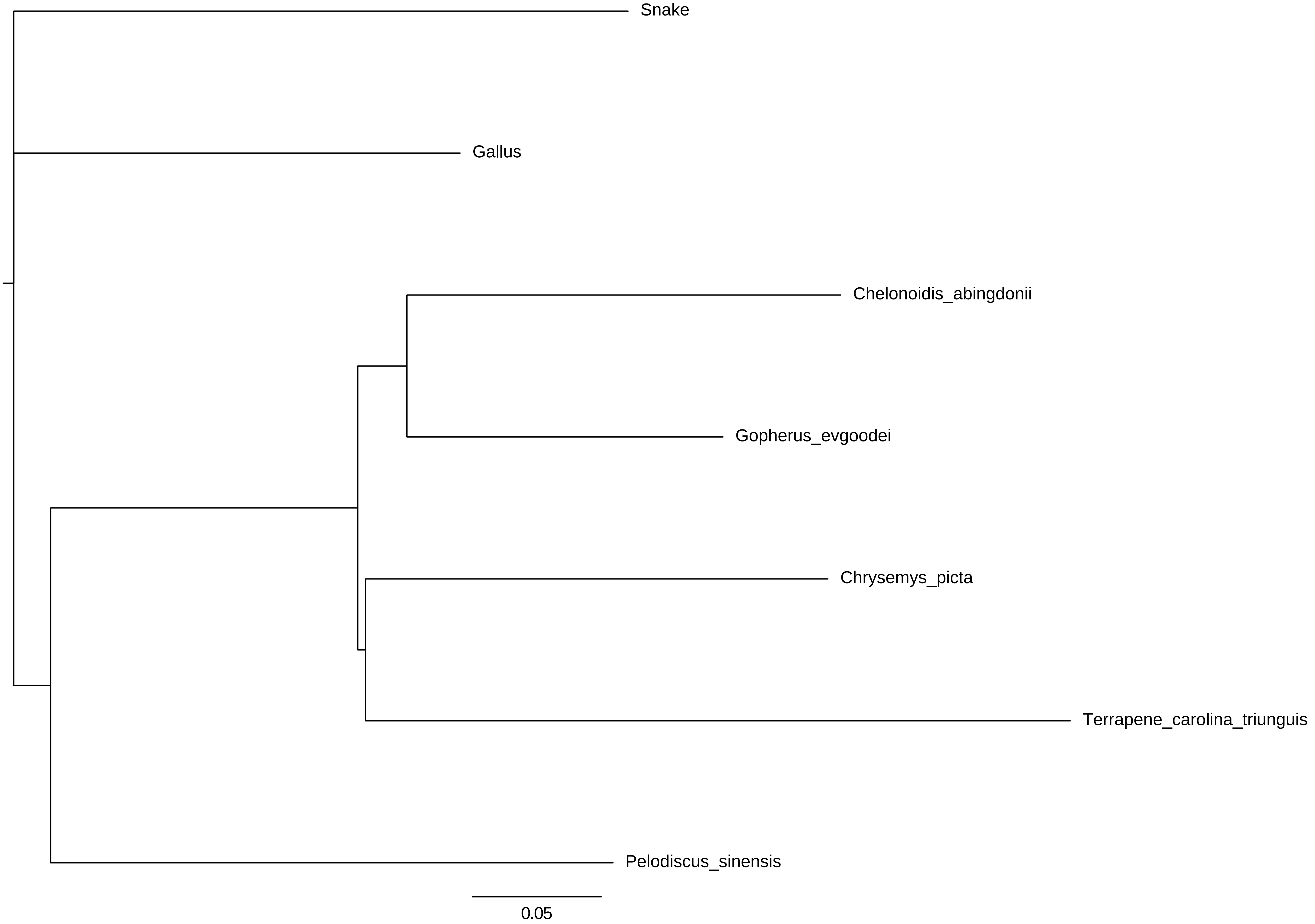

### Figure 4 ΓÇô Figure Supplement 1. Tunicamycin Timecourse.jpg

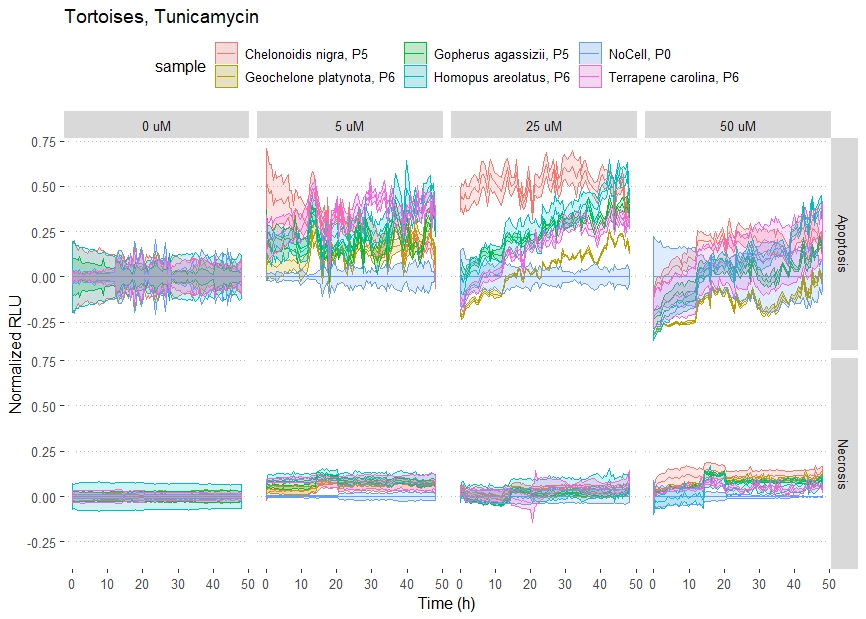

### Figure 4 ΓÇô Figure Supplement 2. Etoposide Timecourse.jpg

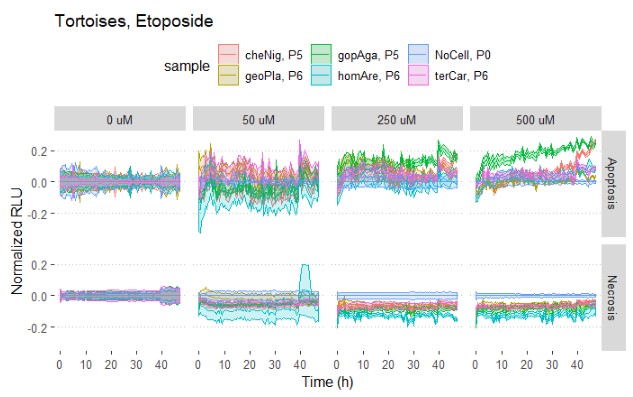

### Figure 4 ΓÇô Figure Supplement 3. Paraquat Timecourse.jpg

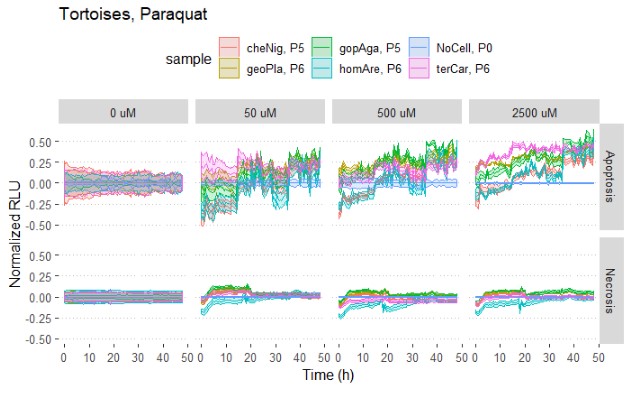

### lifespan-PGLS-Testudines_noGuntheri_realvspred.pdf

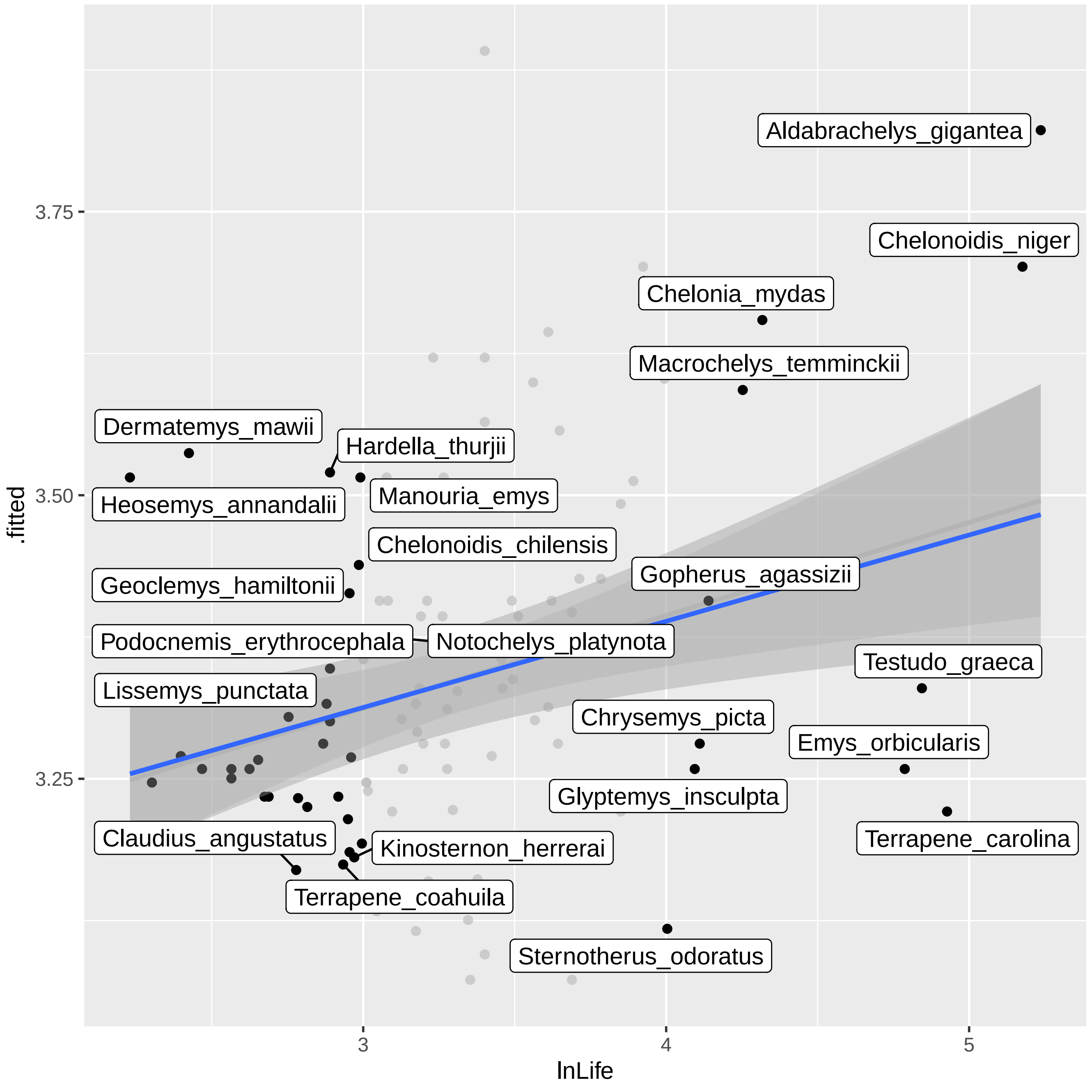

### lifespan-PGLS-Testudines_noGuntheri_residualvsfitted.pdf

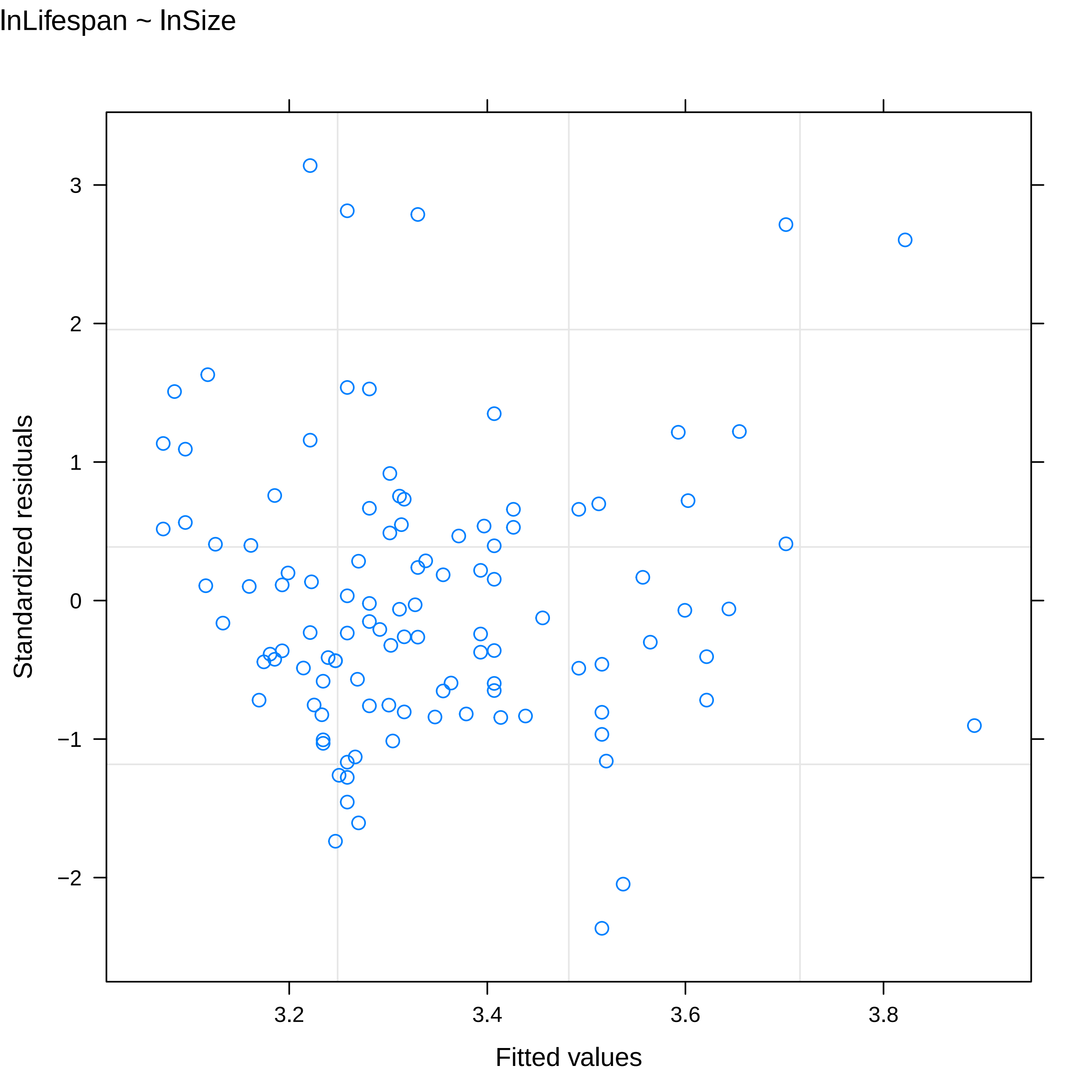

### Testudines_noGuntheri-bodysize.pdf

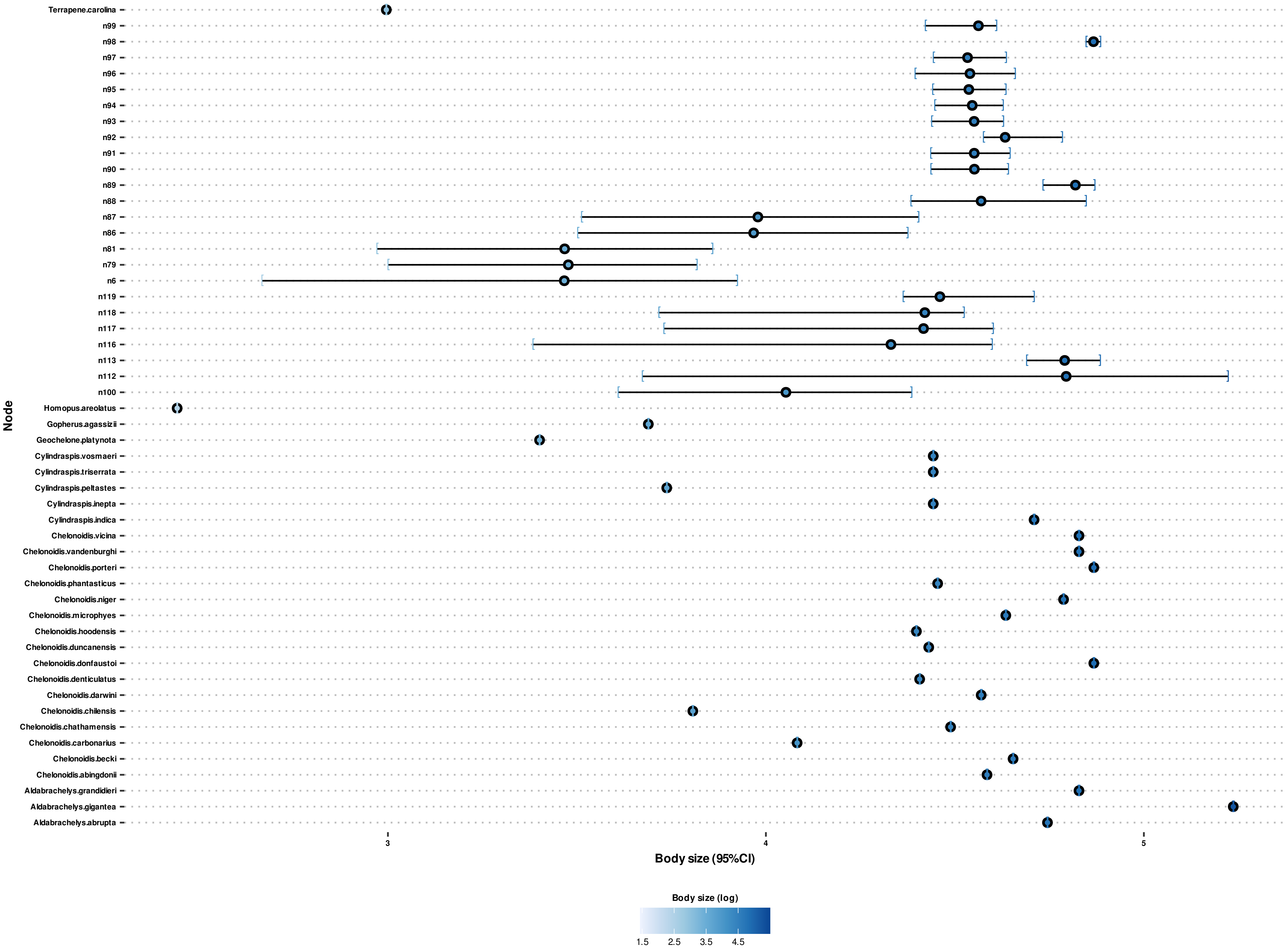

### Testudines_noGuntheri-doubletree.pdf

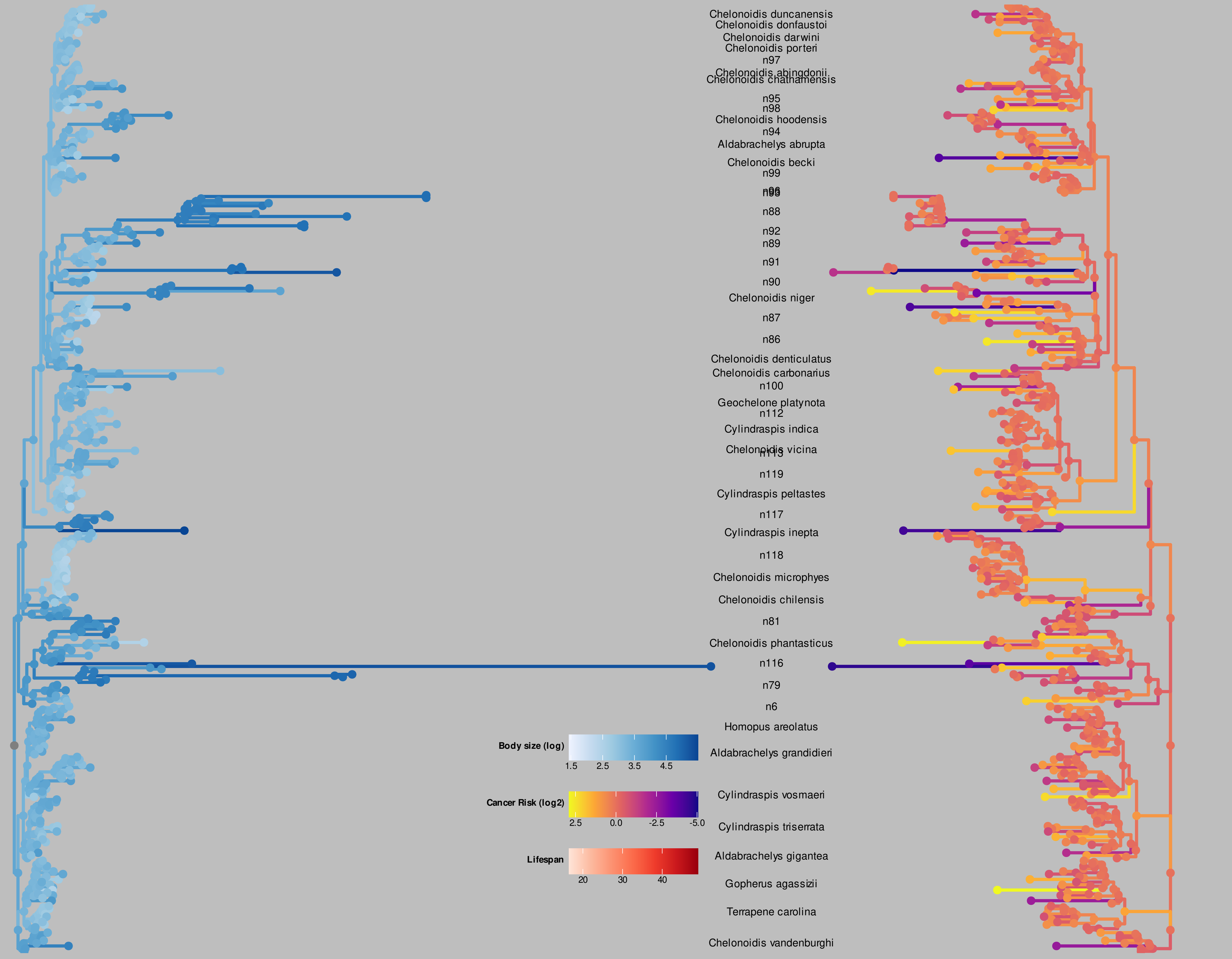

### Testudines_noGuntheri-lifespan.pdf

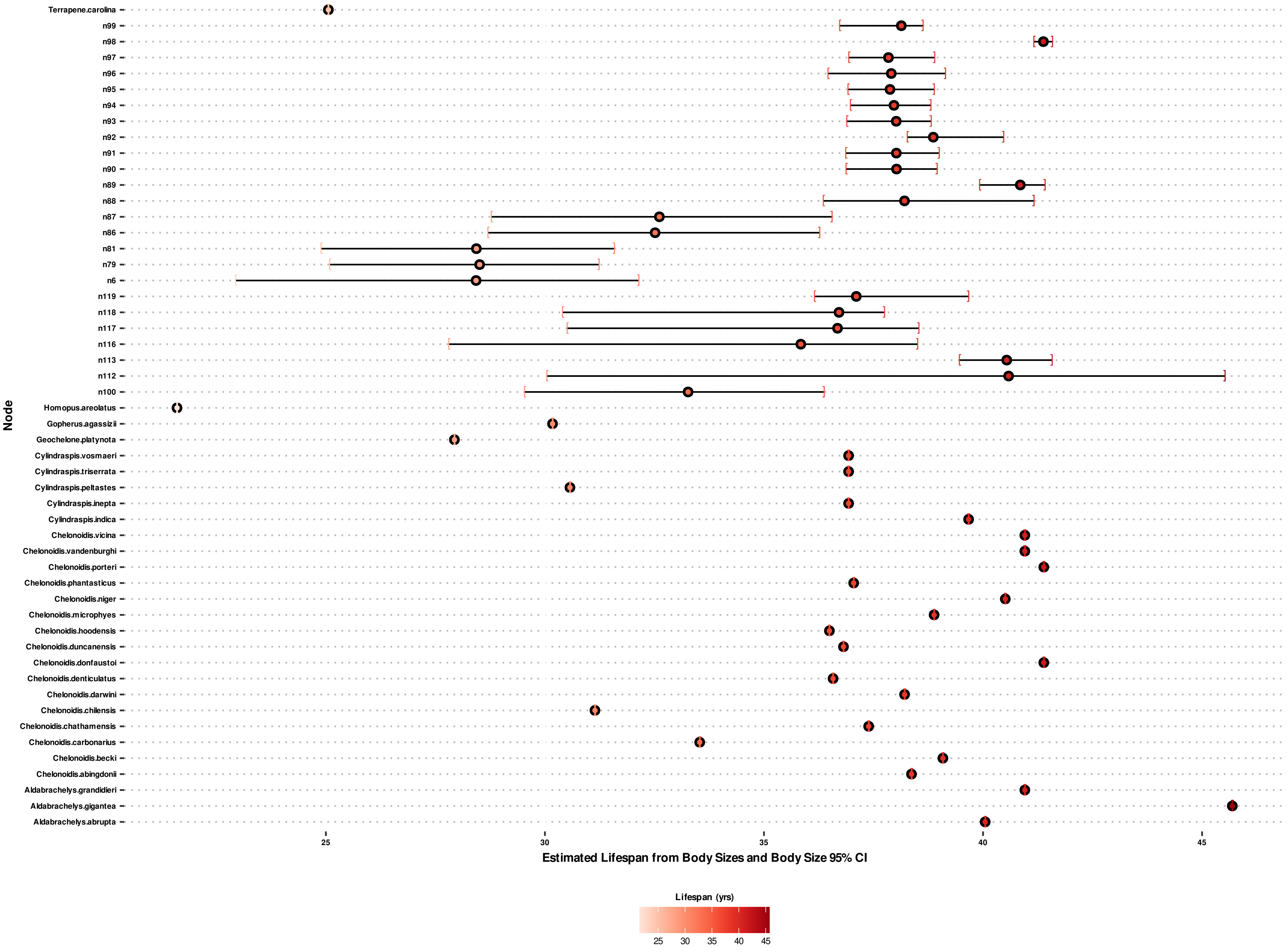

### Testudines_noGuntheri-RICR.pdf

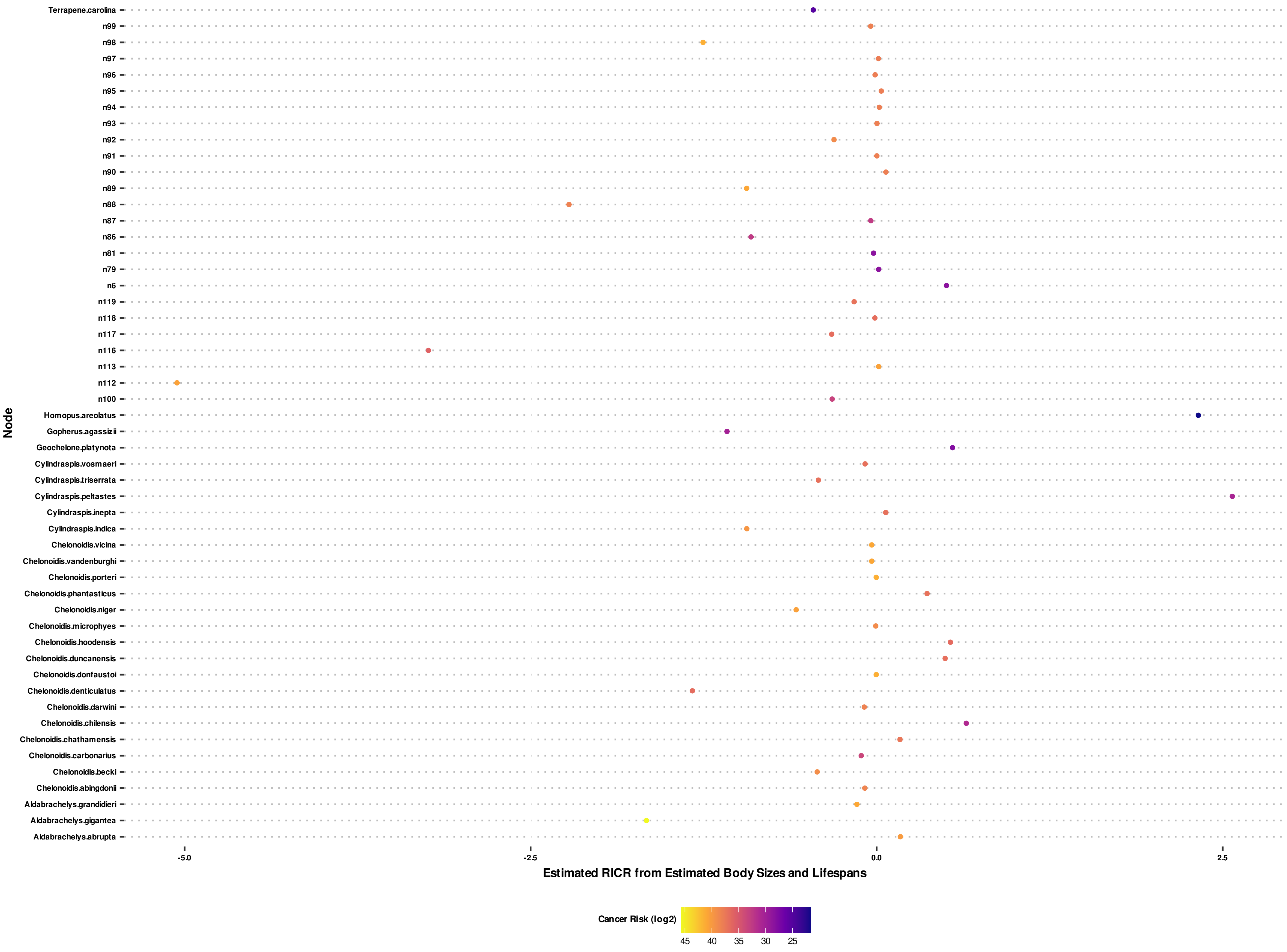
